# EPAS1 Attenuates Atherosclerosis Initiation at Disturbed Flow Sites through Endothelial Fatty Acid Uptake

**DOI:** 10.1101/2023.12.05.570309

**Authors:** Daniela Pirri, Siyu Tian, Blanca Tardajos-Ayllon, Sophie Irving, Francesco Donati, Scott P. Allen, Tadanori Mammoto, Gemma Vilahur, Clint Miller, Torsten Schenkel, Emily V. Chambers, Mark Dunning, Jovana Serbanovic-Canic, Francesco Botrè, Akiko Mammoto, Suowen Xu, Elena Osto, Weiping Han, Maria Fragiadaki, Paul C. Evans

## Abstract

**Background:** Atherosclerotic plaques form unevenly due to disturbed blood flow, causing localized endothelial cell (EC) dysfunction. Obesity exacerbates this process, but the underlying molecular mechanisms are unclear. The transcription factor EPAS1 (HIF2A) has regulatory roles in endothelium, but its involvement in atherosclerosis remains unexplored. This study investigates the potential interplay between EPAS1, obesity, and atherosclerosis.

**Methods:** Responses to shear stress were analysed using cultured porcine aortic EC exposed to flow *in vitro* coupled with metabolic and molecular analyses, and by en face immunostaining of murine aortic EC exposed to disturbed flow *in vivo*. Obesity and dyslipidemia were induced in mice via exposure to high-fat diet or through Leptin gene deletion. The role of Epas1 in atherosclerosis was evaluated by inducible endothelial Epas1 deletion, followed by hypercholesterolemia induction (AAV-PCSK9; high-fat diet).

**Results:** *En face* staining revealed EPAS1 enrichment at sites of disturbed blood flow that are prone to atherosclerosis initiation. Obese mice exhibited substantial reduction in endothelial EPAS1 expression, correlating with hyperlipidaemia. Sulforaphane, a compound with known atheroprotective effects, restored EPAS1 expression and concurrently reduced plasma triglyceride levels in obese mice. Consistently, triglyceride derivatives (free fatty acids) suppressed EPAS1 in cultured EC by upregulating the negative regulator PHD3. Clinical observations revealed that reduced plasma EPAS1 correlated with increased endothelial PHD3 in obese individuals. Functionally, endothelial EPAS1 deletion increased lesion formation in hypercholesterolemic mice, indicating an atheroprotective function. Mechanistic insights revealed that EPAS1 protects arteries by maintaining endothelial proliferation by positively regulating CD36 and LIPG expression to increase fatty acid beta-oxidation.

**Conclusions:** Endothelial EPAS1 attenuates atherosclerosis at sites of disturbed flow by maintaining EC proliferative via fatty acid uptake and metabolism. This endothelial repair pathway is inhibited in obesity, suggesting a novel triglyceride-PHD3 modulation pathway suppressing EPAS1 expression. These findings have implications for therapeutic strategies addressing vascular dysfunction in obesity.

## INTRODUCTION

Atherosclerotic plaques exhibit uneven development, accumulating at arterial regions exposed to disturbed blood flow. These hemodynamic conditions generate mechanical wall shear stress with low magnitude and oscillations in direction that induce localised endothelial cell (EC) dysfunction which drives plaque initiation. The global obesity epidemic is associated with metabolic abnormalities such as dyslipidemia and hyperglycemia that are drivers of atherosclerosis. Disturbed flow enhances the sensitivity of EC to dyslipidemia and hyperglycemia to drive endothelial dysfunction^1,2^, however the molecular mechanisms underlying this link remain poorly understood.

The preservation of endothelial homeostasis relies on an intricate interplay of transcription factors regulating metabolic equilibrium^3^. Notably, the hypoxia-inducible factor (HIF) transcription factors, HIF1A and EPAS1 (or HIF2A), play a central role in this regulatory network^4,5^. Under normal oxygen conditions, these proteins undergo oxygen-dependent proteasomal degradation^6^. However, they accumulate in response to ischemia-induced hypoxia and inflammation, which disrupt the degradation process^7^. HIF1A has a well-established role in angiogenesis^8^. It also accumulates in arteries at sites of disturbed flow where it promotes atherosclerosis through glycolysis and inflammation^9–11^. The function of EPAS1 in vascular biology is less explored. However, recent research suggests its contribution to stabilizing microvascular networks^12–14^ and to physiological processes in several organs^15–21^. However, the potential role of EPAS1 in arterial homeostasis, EC mechanical responses, and atherosclerosis development remains unclear.

Here, we investigate the role of EPAS1 in atherosclerosis development, particularly at disturbed flow sites. We demonstrate that EPAS1 is enriched in these regions, where it limits atherosclerosis by inducing CD36 and LIPG to promote free fatty acid metabolism and enhance endothelial homeostasis. Importantly, our findings reveal that obesity downregulates EPAS1 expression in atheroprone areas, establishing EPAS1 as a crucial intermediate link between obesity and endothelial responses to shear stress. Our research sheds light on the intricate molecular mechanisms connecting obesity, disturbed flow, and atherosclerosis, potentially opening new avenues for further exploration and potential therapeutic interventions.

## MATERIALS AND METHODS

### Mice

Mice with inducible deletion of *Epas1* in EC were generated by crossing *Epas1^fl/fl^* ^22^ mice with *CDH5^Cre-ERT^*^2^ mice^23^. PCR primers used for genotyping are shown in Table S1. To activate Cre, tamoxifen (Sigma) in corn oil was administered intraperitoneally (IP) for 5 consecutive days (2 mg/mouse/d). Two weeks after the first injection of tamoxifen, hypercholesterolemia was induced by intraperitoneal (I.P.) injection of adeno-associated virus containing a gain-of-function mutated version of proprotein convertase subtilisin/kexin type 9 (rAAV8-D377Y-mPCSK9) gene (Vector Core, North Carolina) followed by a HFD (SDS UK, 829100) for 8 weeks as previously described^24^. Some mice were treated with sulforaphane (SFN; Cayman Chemicals) by I.P injection.

*Lep^ob/ob^* mice were obtained from The Jackson laboratory (strain:#00632) and maintained using a standard rodent chow diet. Obesity was induced in C57BL/6N mice by exposing them *ad libitum* to a western diet containing 60% of fat (D12492i, Research Diets) for 25 weeks as described previously^25^. Blood pressure measurements were made using plethismographer CODA (Kent Scientific).

Hyperglycaemia was induced in C57BL/6J mice using Streptozotocin, hereafter referred to as STZ (S1030, Sigma). Mice aged 20 weeks were injected with STZ (150 mg/kg) intraperitoneally and then analysed at 22 weeks.

C57BL/6 and transgenic mice were housed under specific-pathogen free conditions. Animal care and experimental procedures were carried out under licenses issued by the UK Home Office and local ethical committee approval was obtained. All animal procedures conformed to the guidelines from Directive 2010/63/EU of the European Parliament on the protection of animals used for scientific purposes and to IACUC guidelines. All experiments involving animals in Singapore were reviewed and approved by the IACUC of A*STAR Biomedical Sciences Institutes. All mice were on a C57BL/6J background unless otherwise stated.

### Shear stress maps

The lumen geometry of the mouse aorta was derived previously^26^. The unsteady Navier-Stokes equations were solved numerically using the Finite Volume Method (FVM) using the specialised in-house haemodynamics solver haemoFOAM (https://github.com/TS-CUBED/haemoFoam). The vessel geometry was modelled as rigid. Blood was modelled as a non-Newtonian Quemada fluid^27^ with an average haematocrit of 45 percent. The maximum Reynolds number based on the inlet diameter is around 500, therefore, the flow was modelled as laminar. The time dependent mean velocity was digitized from US Doppler images. The aortic root cross section was extruded by 10 diameters to allow for development of the flow and prescription of a Dirichlet inlet boundary condition for the normal velocity as a plug profile at the extruded section. At the exit of the aorta descendent and the branches, a Neumann outlet boundary condition was used with a flow split of 70, 16, 8 and 6 percent for descendent aorta, innominate, common carotid and subclavian, respectively. Numerical solutions were obtained using second order spatial and second order implicit temporal discretization schemes. The mesh size was 0.5 million cells and the time step size was chosen as 1/800^th^ of the cycle time. From the time-dependent WSS, the time-averaged WSS and the oscillatory shear stress index (OSI) at the wall were calculated:

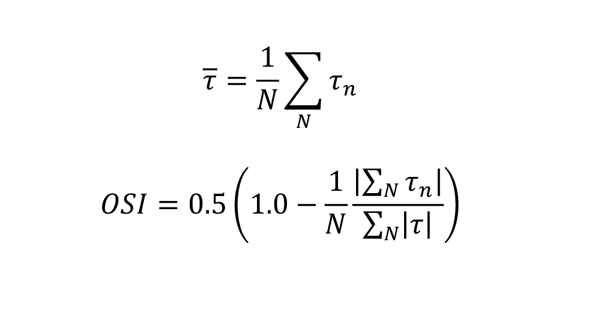

### Endothelial RNA extraction

RNA was extracted from aortic EC using the Qiazol flushing method as described^28^.

### Atherosclerosis plaque analysis

Mice were killed using pentobarbital and perfusion-fixed with PBS followed by 4% PFA. The aorta was dissected, gently cleaned of adventitial tissue, and stained with Oil Red O (Sigma). The surface lesion area was analysed using NIS elements analysis software (Nikon, NY). For analysis of aortic root sections, the upper portion of the hearts were dissected horizontally at the level of the atria and placed in 30% sucrose for 24 h before embedding in paraffin. Serial 7 µm sections were processed for staining with Mayer’s haematoxylin and eosin. ImageJ software was used to calculate the lesion area as a percentage of the total aortic root area.

### Plasma lipid and glucose measurements

Blood samples were collected by terminal cardiac puncture and plasma was separated by centrifugation and analysed using a colorimetric assay (ab65390, Abcam) for detection of total plasma cholesterol and non-HDL cholesterol. Blood glucose and triglycerides levels in venous blood were measured using Accucheck glucometer (Roche Diagnostic) and Accutrend (COBAS, Roche diagnostic), respectively.

### *En face* staining of murine endothelium

The expression levels of specific proteins were assessed in EC at regions of the inner curvature (LOSS site) and outer curvature (HSS site) of murine aortae by *en face* staining. Animals were killed by I.P. injection of pentobarbital and aortae were perfused *in situ* with PBS and then perfusion-fixed with 4% Paraformaldehyde (PFA) prior to harvesting. Fixed aortae were tested by immunostaining using specific primary antibodies. EC were identified by co-staining using anti-CD31 or VE-cadherin (CDH5) antibodies. Nuclei were identified using TO-PRO-3. Stained vessels were mounted prior to visualization of endothelial surfaces *en face* using confocal microscopy (Olympus SZ1000 confocal inverted microscope). The expression of particular proteins at each site was assessed by quantification of the mean fluorescence intensities with standard error of the mean or analysed using ImageJ software (1.49p) to calculate the frequency of positive cells.

### scRNAseq

scRNAseq libraries were generated from age and sex-matched *Epas1^EC-KO^*and *Epas1^EC-WT^* mice using the SORT-seq protocol as described previously^29^. Dimensional reduction and clustering of our scRNA-seq dataset were performed using the R package Seurat (v3.1). Differential expression analysis was performed between clusters, and marker genes for each cluster were determined with the Wilcoxon rank-sum test with p value<0.001 and a minimum log-fold change threshold of 0.25 using Seurat and BBrowser (version 2.10.10). Differential expression analysis, heatmaps of gene expression embedded on hierarchical clustering, tSNE representations showing the expression of defined gene sets/ GO pathways and expression of single transcripts on the tSNE embedding were performed using the software BBrowser (version 2.10.10). tSNE representation of the expression of defined GO pathways shows the signature score, which indicates the sum of all features in each GO pathway.

### PAEC culture and exposure to WSS

Pig aortae from 4-6 months old animals were obtained immediately after slaughter from a local abattoir. EC were harvested using collagenase (1 mg/ml for 10 minutes at room temperature) and were cultured using M199 (Gibco) media. Experiments were performed using cells from multiple donors that were not pooled. PAEC at passage 3-5 were seeded onto gelatin-coated Ibidi µ-Slides I^0.4^ (Luer ibiTreat, ibidi^TM^) and used when fully confluent. Flowing medium was then applied using the Ibidi pump system to generate HSS (13 dyn/cm^2^), LSS (4 dyn/cm^2^) or low oscillatory WSS. For low oscillatory WSS, PAEC were exposed to a repeated cycle of 2 h of oscillatory flow (+/-4 dyn/cm^2^, 0.5 Hz), followed by 10 min of unidirectional flow (+4 dyn/cm^2^), to ensure redistribution of nutrients^30^. The slides and pump apparatus were placed in a cell culture incubator at 37°C. Alternatively, PAECs were cultured in a 6 well plate and exposed to shear using the orbital system as described^9^. Some sheared cultures were also exposed to FFAs (oleic acid; 0.25 mM).

### Gene silencing

PAEC cultures were transfected with a pool of shRNA sequences targeting porcine *EPAS1* (V3SVHSHC, Dharmacon) using the Dharmacon™SMARTvector Lentiviral system following the manufacturer’s instructions. Non-targeting lentiviral shRNA sequence was used as a control (V16060303, Dharmacon). All lentiviral transductions were performed at 30 multiplicity of infection in the presence of 6µg/ml of DEAE-dextran hydrochloride (Sigma).

### Real time PCR

RNA was extracted using the RNeasy Mini Kit (74104, Qiagen) and reverse transcribed into cDNA using the iScript cDNA synthesis kit (1708891, Bio-Rad). QRT-PCR was used to assess the levels of transcripts with gene-specific primers (Table 1). Reactions were prepared using SsoAdvanced universal SYBR®Green supermix (172-5271, Bio-rad) and following the manufacturer’s instructions, and were performed in triplicate. Expression values were normalized against the house-keeping genes *B2M* (pig), Hprt (human, mouse). Data were pooled from at least three independent donors and mean values were calculated with SEM.

**Table1.**
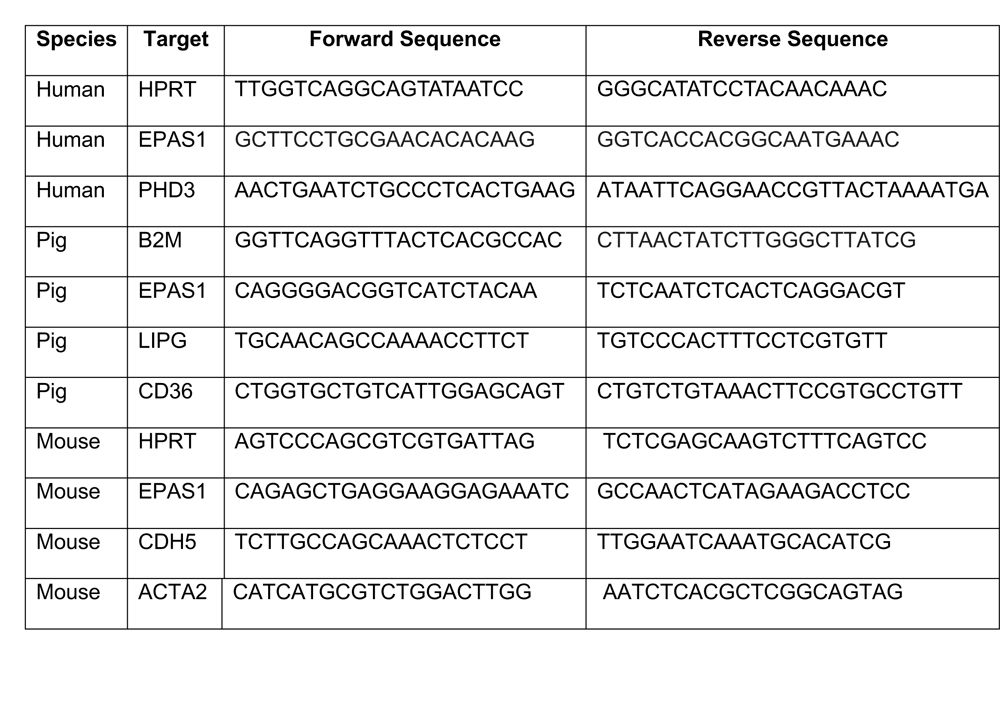

### XF24 metabolic fuel flex assay

PAEC were plated in a Cell Tak-coated XF24 cell culture plate (Agilent) in XF assay media pH 7.4 (Agilent) supplemented with 2 mM glutamine and 25 mM glucose. OCR was measured for 32 min prior to the addition of Etomoxir (6 µM) to analyse the contribution of fatty acid β -oxidation to OCR, then the addition of BPTES (1µM) and UK5099 (6 µM) to analyse the contribution of glycolysis and glutamine to OCR. The contribution of fatty acid β-oxidation to OCR (fatty acid dependency) was calculated as follows: [OCR prior to Etomoxir - OCR after Etomoxir] / [OCR prior to Etomoxir - OCR after BPETS+UK5099].

### Immunofluorescent staining of cultured EC

PAEC were fixed with PFA (4%) and permeabilised with Triton X-100 (0.1%). Following blocking with goat serum for 30 min, monolayers were incubated for 16 h with primary antibody against Ki67 and AlexaFluor488. Nuclei were identified using DAPI (Sigma). Images were taken with a widefield fluorescence microscope (LeicaDMI4000B) and analysed using ImageJ software (1.49p) to calculate the frequency of positive cells. Isotype controls or omission of the primary antibody was used to control for non-specific staining.

### Immunoblotting

Total cell lysates were isolated using lysis buffer (containing 2% SDS, 10% Glycerol and 5% β-mercaptoethanol). Primary antibodies used and concentrations are described in Table S2. HRP-conjugated secondary antibodies (Dako) and chemiluminescent detection was carried out using ECL Prime^®^ (GE Healthcare). Membranes were imaged using the Gel Doc XR+ system (Biorad).

### Clinical adipose and plasma samples

Human subcutaneous adipose tissues (n = 29) were collected from obese (BMI>30) and non-obese (BMI<30) individuals. These tissues were obtained as surgical discards from patients undergoing abdominal surgeries. Following surgical removal, the samples were promptly placed in ice-cold HEPES buffer prior to EC isolation as described^31^. De-identified demographic data of the patients were retrieved using the Generic Clinical Research Database at the Medical College of Wisconsin. All procedures were approved by the Institutional Review Board of MCW and Froedtert Hospital. Peripheral blood samples were taken from severe obese subjects with BMI>38 (n=5) and from healthy non-obese subjects (n=24). The local Research and Ethic Committees in Zurich, Switzerland approved the study (Ethic Nr.KEK-ZH_Nr.2013-0389). Serum samples were generated immediately and stored at −80°C until analysis. EPAS1 levels in serum were quantified by ELISA using a commercial kit (CUSABIO, CSB-E12113h). All patients gave written informed consent. Studies were performed according to the principles of the Declaration of Helsinki.

### Statistical analysis

Data are presented as mean values ± SEM. Statistical analyses were performed by ANOVA or t-tests as stated using GraphPad Prism software. The degree of significance is as following: *p <0.05; **p<0.01; ***p< 0.001. The test performed is indicated in the figure legend.

## RESULTS

### EPAS1 is enriched at an atheroprone site exposed to low oscillatory shear stress

To analyse whether EPAS1 is responsive to flow, we exposed porcine aortic EC (PAEC) to varying shear stress conditions using a parallel plate system. Immunoblotting revealed that EPAS1 expression was increased in PAEC exposed to low and oscillatory shear stress (LOSS; 4 dynes/cm2; bidirectional with 1Hz frequency) compared to low shear stress (LSS; 4 dynes/cm2) or unidirectional high shear stress (HSS; 13 dynes/cm2) (Fig. 1A). We then measured EPAS1 levels in EC of the mouse aortic arch which is known to generate disturbed flow associated with atherosclerosis initiation^32^. CT-angiography coupled to computational fluid dynamics showed that the inner curvature was exposed to relatively low time-averaged WSS magnitude and more oscillations in direction (i.e. LOSS) compared to the outer curvature (Fig. 1B), consistent with prior analyses^26,32^. In agreement with our in-vitro findings, *en face* immunofluorescent staining of the murine aorta revealed that EPAS1 was significantly enriched at the LOSS region (inner curvature) compared to HSS regions (Fig. 1C). A proportion of EPAS1 at the LOSS site localized to the nucleus suggesting that it is active (Fig. 1C).

**Figure 1.**
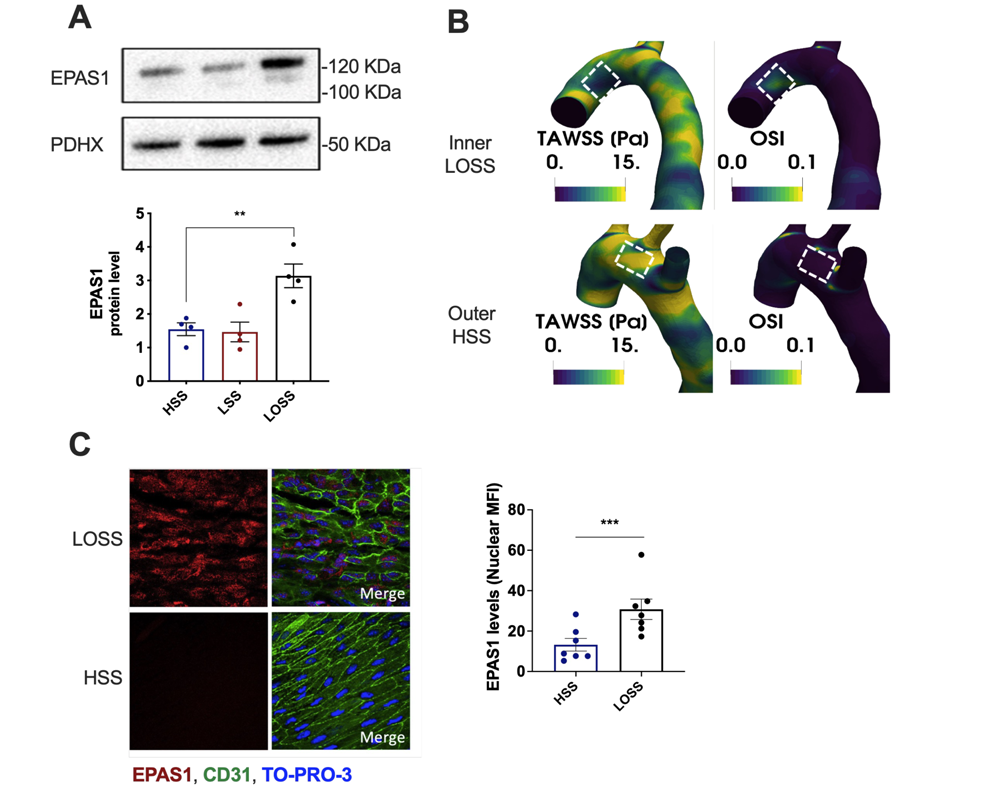
EPAS1 is enriched at an atheroprone site exposed to low and oscillatory shear stress. (A) PAEC were seeded on μ-slides and cultured under HSS, LSS or LOSS for 72h using the Ibidi system. Protein levels of EPAS1 were quantified by immunoblotting. Representative images and mean values normalized to the level of PDHX (n=4) are shown. Differences between means were analysed using a one-way ANOVA. (B) Time-averaged WSS (TAWSS) and oscillatory shear index (OSI) were mapped onto the geometry of the murine aortic arch. Representative images are shown and regions exposed to LOSS (inner) or HSS (outer) are marked. (C) Aortic arches were isolated from C57BL/6 mice aged 6-8 weeks and *en face* immunostaining was performed using anti-EPAS1 antibodies (red). Endothelium was co-stained (EC; green) and nuclei detected using TO-PRO-3 (blue). EPAS1 levels were quantified at LOSS and HSS regions. Each data point represents an animal. Differences between means were analysed using a t-test.

Thus, EPAS1 is enhanced by LOSS *in vitro* and it is enriched at an atheroprone region of the murine aorta exposed to LOSS.

### Endothelial EPAS1 levels are regulated by metabolic status

Dysregulation of the HIF pathway has been associated with the development of several cardiovascular risk factors such as obesity and diabetes type 2^33,34^. Nevertheless, the possible effect of these risk factors on endothelial HIF expression has not been explored. To assess the effect of obesity on aortic endothelial EPAS1 levels, we exposed wild type mice to high fat diet (HFD) or normal chow diet for 22 weeks. HFD mice were significantly heavier than chow diet mice (Fig 2A). Weight gain was also associated with hyperglycaemia (Fig 2B), increased triglycerides levels (Fig 2C) and raised total cholesterol levels (Fig 2D). We further tested for HDL cholesterol which showed no differences between chow and HFD mice, while LDL/VLDL ratio was significantly increased in HFD mice (Fig S1A). In addition, HFD did not elicit significant changes of the mouse systolic blood pressure despite elevated diastolic pressure (Fig S1B). *En face* staining of the mice aorta revealed a dramatic decrease in EPAS1 levels at the LOSS region in mice exposed to HFD compared to chow fed control mice (Fig 2E). Our data suggest that HFD feeding associated with obesity reduces EPAS1 expression at a LOSS atheroprone region.

**Figure 2.**
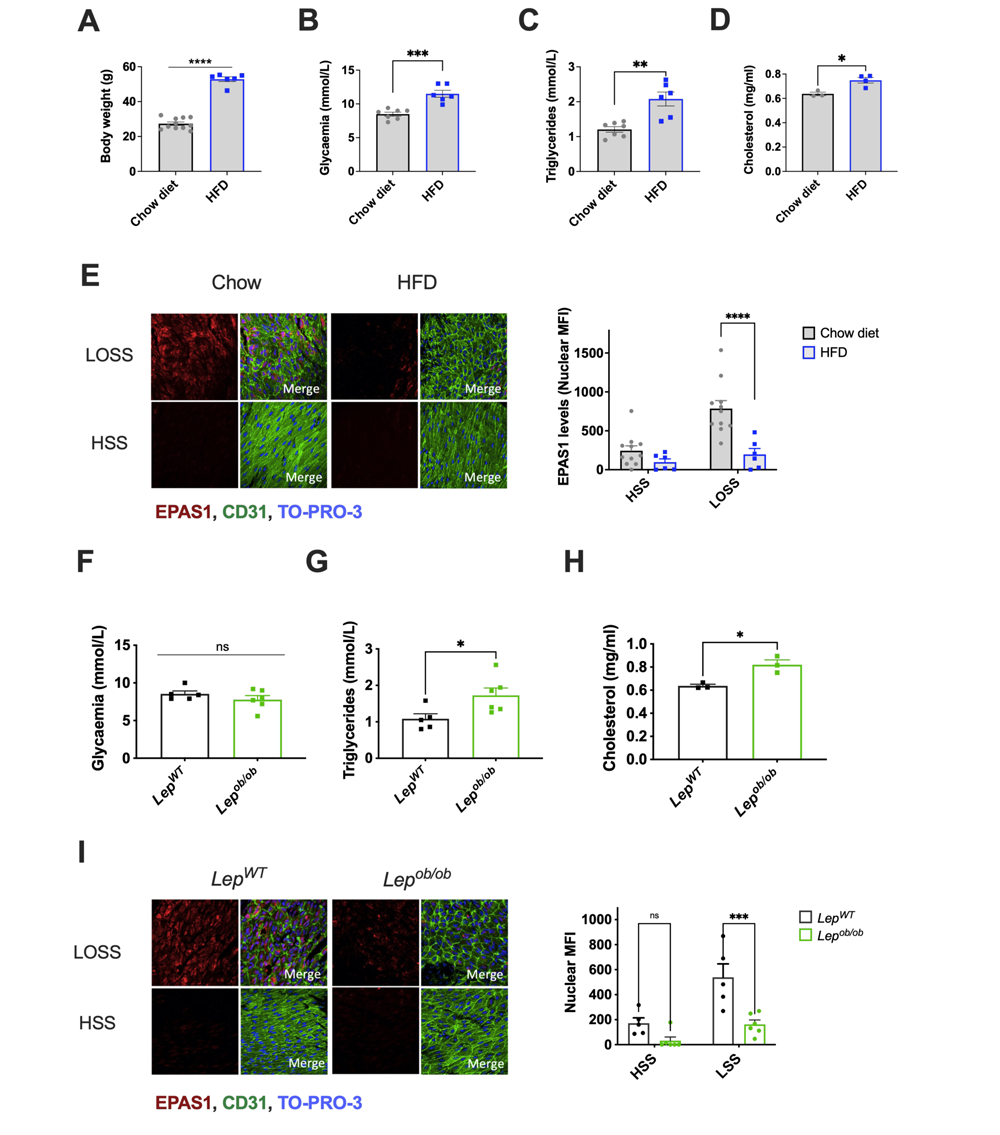
High fat feeding and obesity reduced EPAS1 levels at atheroprone aortic endothelium. (A-E) C57BL/6N mice aged 5 weeks were exposed to HFD or to standard chow for 25 weeks. Body weight (A), glycaemia (B), plasma triglycerides (C) and plasma cholesterol (D) were measured. (E) Aortic EC were stained *en face* using anti-EPAS1 antibodies (red) and fluorescence was quantified at LOSS and HSS regions. (F-I) *Lep^ob/ob^* mice and littermate controls (*Lep^wt/wt^*) aged 22 weeks were analysed. Glycaemia (F), plasma triglycerides (G) and plasma cholesterol (H) were measured. (I) Aortic EC were stained *en face* using anti-EPAS1 antibodies (red) and fluorescence was quantified at LOSS and HSS regions. Endothelium was co-stained (EC; green) and nuclei detected using TO-PRO-3 (blue).

To confirm the influence of obesity on endothelial EPAS1 levels, we analysed an alternative model of obesity using mice with homozygous deficiency in leptin (*Lep^ob/ob^*) which gained weight more rapidly than littermate controls (Fig. S1C). Heterozygous mice (*Lep^ob/+^*) did not exhibit accelerated weight gain and were therefore excluded from subsequent analysis (Fig. S1C). *Lep^ob/ob^* mice were characterised by several metabolic changes including transient hyperglycaemia, raised within 14 weeks (Fig. S1D) which normalised at later time points of analysis (Fig. 2F). *Lep^ob/ob^* mice exhibited increased levels of plasma TG (Fig. 2G), total cholesterol (Fig 2H), and increased LDL/VLDL ratio with no changes in HDL cholesterol levels (Fig S1E). It was observed that EPAS1 levels at LOSS regions were strikingly reduced in *Lep^ob/ob^* mice compared to wild-type controls (Fig. 2I). Therefore, obesity induced through either exposure to a HFD or genetic loss of leptin led to a substantial reduction in EPAS1 levels in aortic endothelium.

We next sought to define the mechanism of EPAS1 reduction in obesity. The potential contribution of hypertension was eliminated because *Lep^ob/ob^* mice exhibited a significant decrease in both systolic and diastolic blood pressure compared to controls (Fig S1F). Despite the severe obesity status, this phenomenon was predictable and previously associated with the *Lep^ob/ob^* phenotype^35^. We therefore focussed on the effects of metabolic alterations in obesity on EPAS1 expression. The potential contribution of hyperglycamia was analysed by treating mice with streptozotocin (STZ) to ablate the pancreas. STZ induced sustained hyperglycaemia in chow fed mice (Fig. 3A) and had no effect on TG levels (Fig. 3B). *En face* staining revealed that EPAS1 expression at the LOSS region of the aorta was unaltered in STZ-treated mice (Fig. 3C), indicating that hyperglycaemia is insufficient to suppress EPAS1. To corroborate this finding, we investigated the levels of EPAS1 in the aorta of *Lep^ob/ob^* mice during their hyperglycaemic phase at 14 weeks. Hyperglycaemic *Lep^ob/ob^* mice did not exhibit altered EPAS1 at the LOSS region (Fig. S2A), confirming that EPAS1 suppression in obesity can be uncoupled from hyperglycaemia.

**Figure 3.**
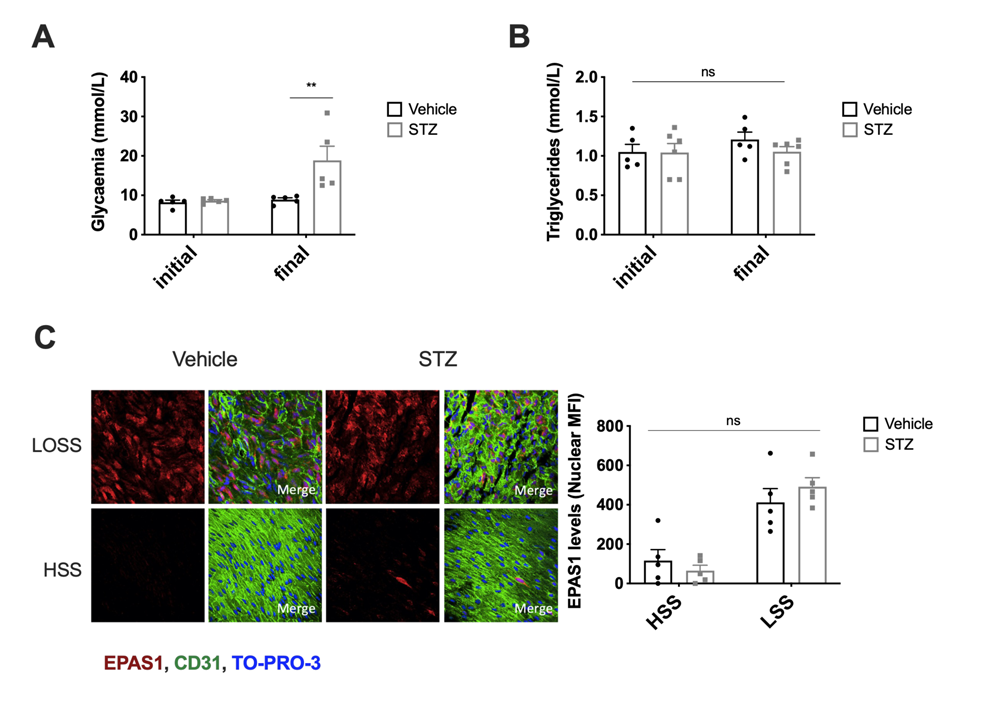
Hyperglycaemia does not contribute to EPAS1 reduction. Mice (C57BL/6J) aged 20 weeks were treated with STZ (Day 0) and analysed two weeks later. Glycaemia (A) and plasma triglycerides (B) were measured. (C) Aortic EC were stained *en face* using anti-EPAS1 antibodies (red) and fluorescence was quantified at LOSS and HSS regions. Endothelium was co-stained (EC; green) and nuclei detected using TO-PRO-3 (blue). Each data point represents an animal. Differences between means were analysed using a two-way ANOVA.

Next the potential role of hyperlipidaemia was analysed by treatment of obese mice with sulforaphane, a compound from broccoli and other cruciferous vegetables that is known to modify lipid profiles^36^ and protect endothelium. Sulforaphane treatment significantly reduced plasma levels of TGs in obese mice exposed to the HFD (Fig. 4A), whereas glycaemia (Fig. 4B), body weight (Fig. S3A), and plasma levels of total cholesterol (Fig. S3B), HDL and LDL/VLDL (Fig. S3C) were unaltered. Sulforaphane treatment of obese mice restored the expression of EPAS1 specifically at the LOSS region of the aorta (Fig. 4C). These data show that obesity reduces the expression of EPAS1 at sites of LOSS, and this can be rescued by the TG-lowering compound sulforaphane. These observations led us to hypothesize that chronic exposure of EC to TGs may be responsible for EPAS1 suppression in obesity. This was tested directly by exposing PAEC cultured under LOSS to free fatty acids (FFAs) which are produced from local hydrolysis of TG. Exposure to FFAs led to a significant reduction in EPAS1 levels (Fig. S4A) and concomitantly increased levels of PHD3 (Fig. S4B), a protein that negatively regulates EPAS1 via proteasomal degradation^22^. These data suggest that EPAS1 reduction in obesity may result from hypertriglyceridemia leading to the induction of endothelial PHD3 by FFAs.

**Figure 4.**
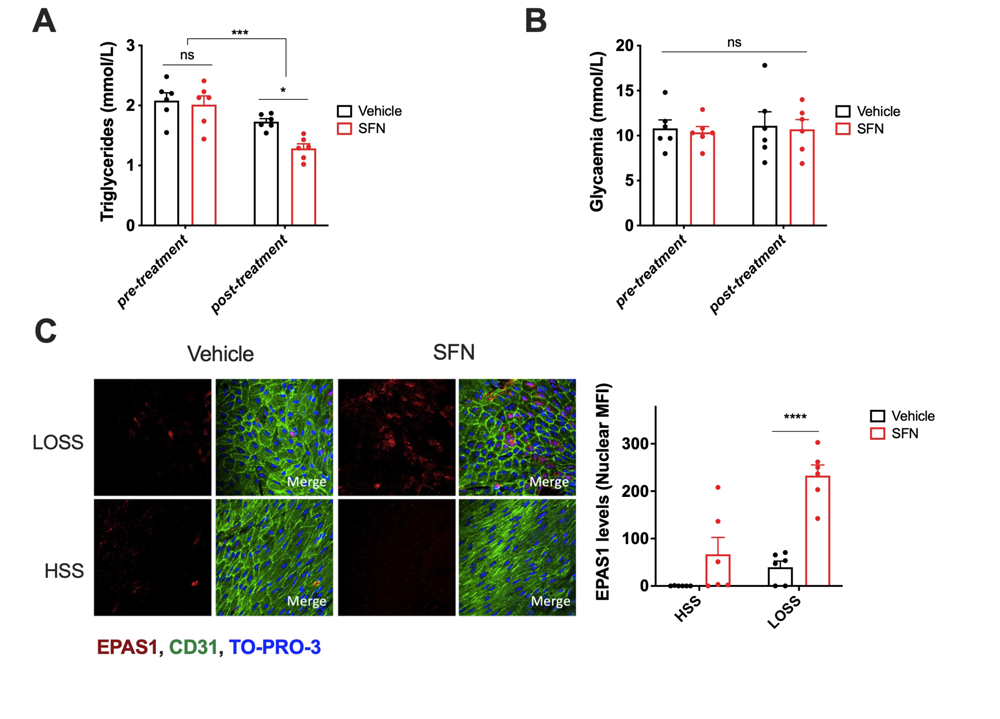
Sulforaphane rescues EPAS1 in obese mice. C57BL/6N mice aged 5 weeks were exposed to HFD for 25 weeks (pretreatment group). They were then treated with sulforaphane (SFN; daily I.P. injections 5 mg/kg for 3 days) or with vehicle for 3 days, with both groups receiving a HFD for that period. Plasma triglycerides (A) and glycaemia (B) were measured pretreatment and in SFN-treated and vehicle control groups. (C) Aortic EC were stained *en face* using anti-EPAS1 antibodies (red) and fluorescence was quantified at LOSS and HSS regions in SFN-treated and control groups. Endothelium was co-stained (EC; green) and nuclei detected using TO-PRO-3 (blue). Each data point represents an animal. Differences between means were analysed using one-way ANOVA (A, B) or two-way ANOVA (C).

The clinical relevance of our findings was assessed by comparing the PHD3-EPAS1 pathway in obese versus non-obese individuals. Since sampling of arterial EC was not feasible, we quantified EPAS1 protein levels in plasma as a surrogate. A cohort of both male and females with no significant difference in age (Fig. S5A) was analysed. ELISA revealed that EPAS1 was detectable in all healthy controls studied but was either absent or significantly reduced in plasma samples from obese individuals (Fig. S5B). This pathway was further interrogated by measuring *EPAS1* and *PHD3* transcripts in microvascular EC isolated from subcutaneous adipose of obese patients and non-obese controls. qRT-PCR revealed that *PHD3* levels were significantly increased in obese individuals compared to controls whereas EPAS1 mRNA expression was unaltered (Fig. S5C). Overall, these data suggest that EPAS1 is suppressed in obese individuals associated with elevated *PHD3* expression in EC, a trend that is consistent with our observations in murine models of obesity.

### *Epas1* protects the endothelium against atherosclerosis by regulating metabolic and proliferative pathways

To analyse the role of Epas1 in atherosclerosis we deleted *Epas1* from the endothelium of mice using *Cdh5^CreERT^*^2^*^/+^* line crossed with a *Epas1^fl/fl^* mouse to generate *Epas1^EC-KO^* and control *Epas1^EC-WT^* mice (Fig. 5A). Tamoxifen treatment of *Epas1^EC-KO^* and control *Epas1^EC-WT^* mice induced deletion of *Epas1* which was validated by qRT-PCR of endothelial RNA (Fig. S6A) and by *en face* staining (Fig. S6B). *Epas1^EC-KO^* and control *Epas1^EC-WT^* mice were treated with AAV-PCSK9 and exposed to a HFD for 8 weeks to generate hypercholesterolemia (Fig. 5A). Endothelial deletion of *Epas1* had no effect on plasma cholesterol levels (Fig. 5B). However, atherosclerotic plaques in the aorta were significantly larger in *Epas1^EC-KO^*mice compared to controls (Fig. 5C). Plaques formation was also measured at the aortic root, confirming the presence of larger plaques in *Epas1^EC-KO^* mice compared to *Epas1^EC-WT^*controls (Fig. 5D). These data show that endothelial *EPAS1* is atheroprotective.

**Figure 5.**
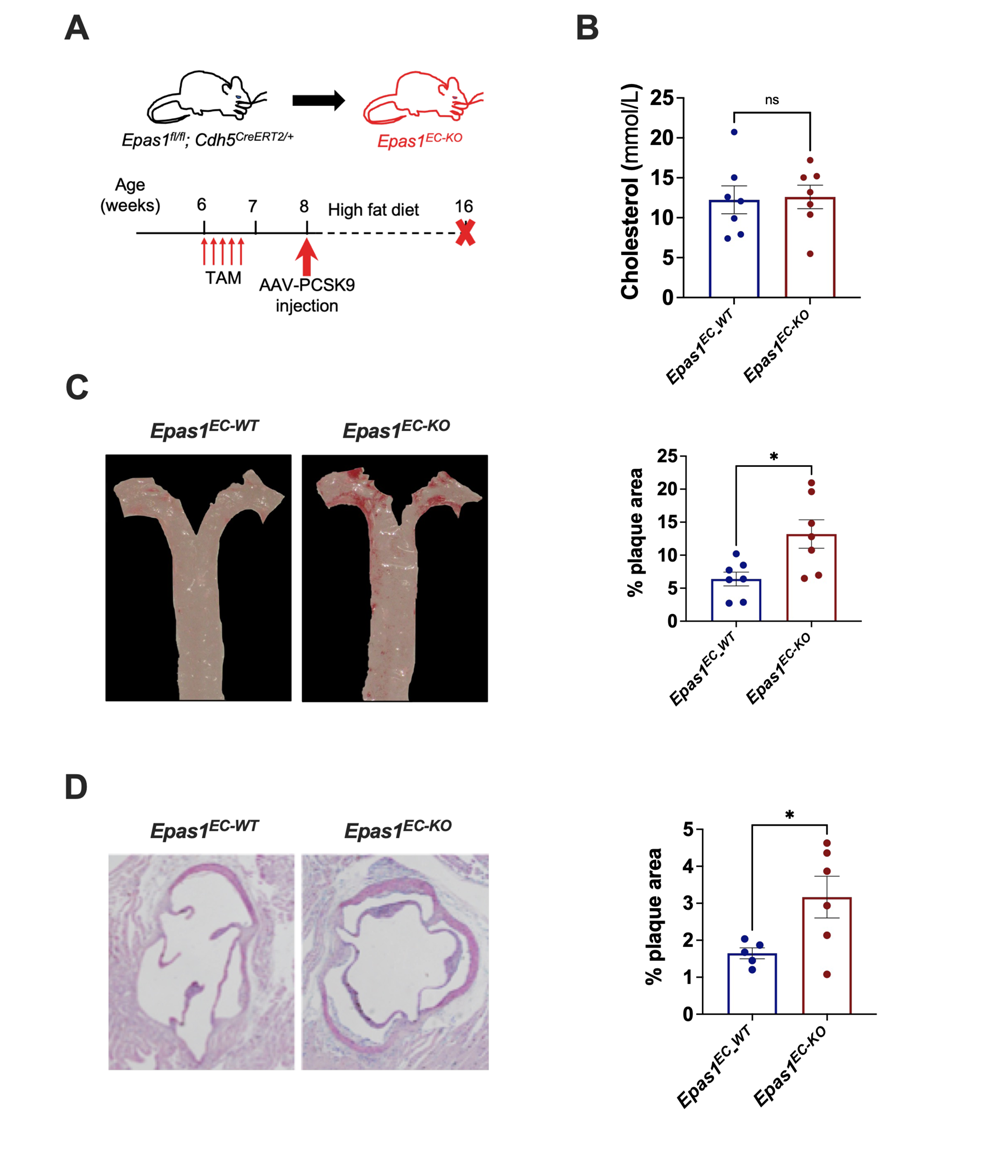
Endothelial *Epas1* protects against atherosclerosis. (A) Timeline of *Epas1* deletion in a model of hypercholesterolemia. *Epas1^EC-KO^* mice aged 6 weeks and *Epas1^EC-WT^* mice received five intraperitoneal injections of tamoxifen and one injection of PCSK9-AAV virus at specified time points. After 8 weeks fed with high fat diet, the mice were culled and plaque area quantified. (B) Plasma cholesterol levels were measured in *Epas1^EC-KO^* mice and *Epas1^EC-WT^* controls. (C) Quantification of plaque burden in the aorta was determined by calculating the percentage of aortic surface area covered by plaque for *Epas1^EC-KO^*mice and *Epas1^EC-WT^* controls. (D) Quantification of plaque burden in the aortic roots of *Epas1^EC-KO^* mice and *Epas1^EC-WT^*controls. Each data points represents one mouse and mean ± SEM are shown. Differences between means were analysed using an unpaired *t*-test.

To elucidate the function of endothelial EPAS1 we performed single cell RNAseq from aortas of *Epas1^EC-KO^* and control (*Epas1^EC-WT^*) mice. Endothelial cells were isolated by enzymatic digestion and CD31^+^ CD45^-^ TO-PRO-3^-^ EC and purified by FACS and processed for scRNAseq (Fig. S7A). Our sort-seq analysis identified 6 distinct clusters (Fig. S7B). Of these, clusters 0-4 consistently expressed multiple EC markers whereas cluster 5 contained VSMC-like features and was subsequently removed from analysis of EC heterogeneity (Fig. S7C). The majority of cells in cluster 0 were from control mice, whereas clusters 1 and 4 contained cells mainly from *Epas1^EC-KO^*mice (Fig. S7D and Fig. S7E), indicating that *Epas1* influences EC heterogeneity. Functional annotation revealed diverse functions associated with *Epas1*-regulated clusters, most notably several GO terms associated with metabolic processes (long-chain fatty acid transport; cGMP-mediated signalling – Lipoprotein; Response to hypoxia) were enriched in cluster 4 (Fig. S7F). These data suggest that *Epas1* may be involved in fatty acid/lipoprotein metabolism in arterial endothelium.

### EPAS1 promotes fatty acid β-oxidation and proliferation under atheroprone shear stress

To investigate directly whether EPAS1 regulates fatty acid metabolism, we investigated the effects of *EPAS1* gene silencing on metabolic processes in cultured PAEC. A lentiviral-small hairpin RNA (shRNA) effectively reduced EPAS1 at both the mRNA (Fig. S8A) and protein levels in PAEC cultured under LOSS conditions (Fig. S8B). Next, the effects of *EPAS1* silencing on oxygen consumption rates (OCR) was assessed by performing a fuel flux assay using a Seahorse metabolic bioanalyser. Here, OCR was measured in PAEC exposed to LOSS that were sequentially treated with Etomoxir (inhibitor of long chain fatty acid oxidation), followed by a combination of UK5099 (inhibitor of glucose oxidation pathway) and BPTES (inhibitor of glutamine oxidation) (Fig. S8C). It was observed that *EPAS1* silencing caused a striking reduction in basal OCR of EC exposed to LOSS (Fig. 6A; Fig. S8C) and that OCR was strongly dependent on fatty acid β-oxidation (Fig. S8D). These data indicate that EPAS1 is a positive regulator of fatty acid metabolism.

**Figure 6.**
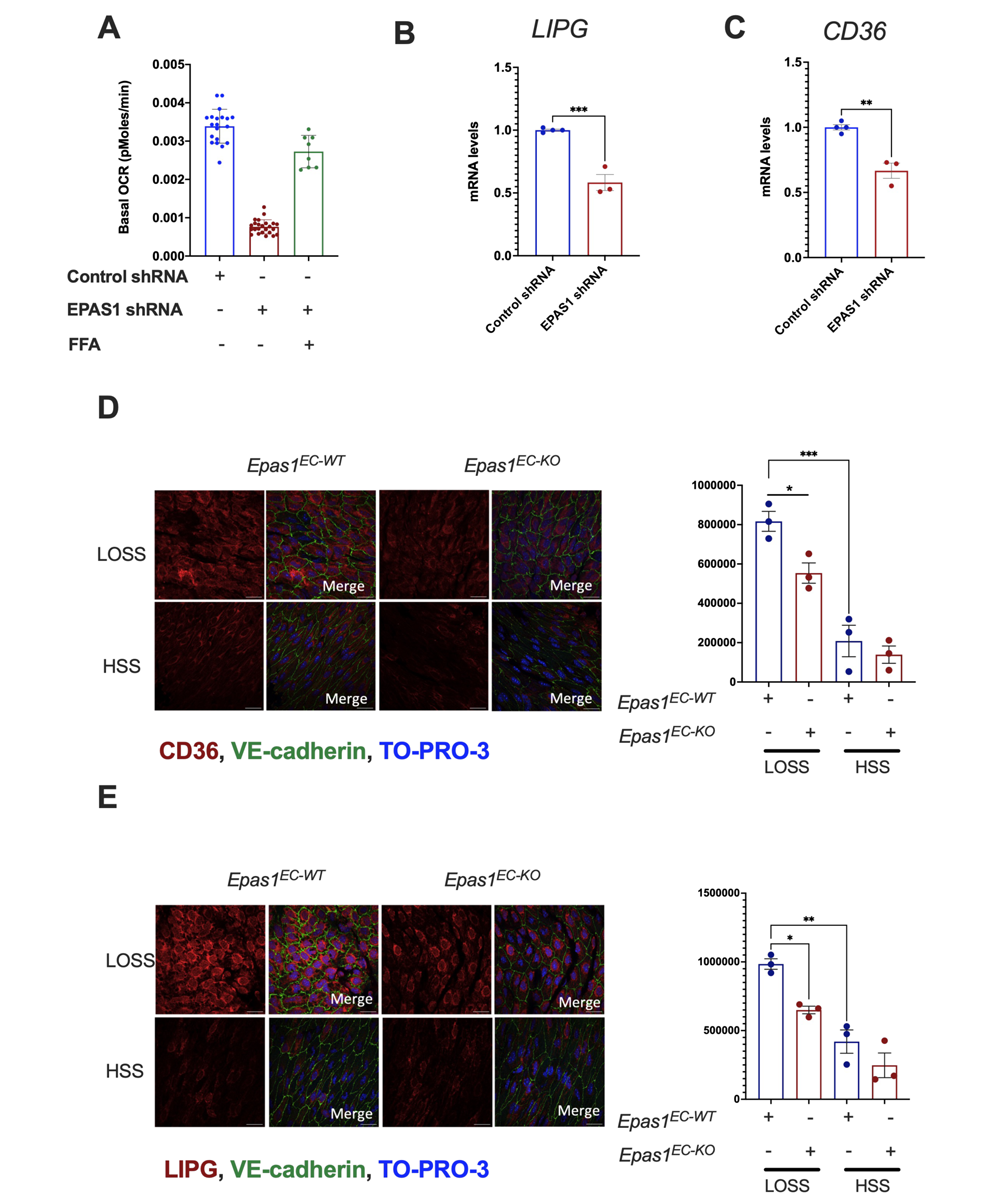
Endothelial EPAS1 controls fatty acid metabolism via LIPG and CD36. (A-C) PAEC were treated with shRNA targeting *EPAS1* or with scrambled control and exposed to LOSS for 72 h using the orbital system. (A) Some cultures were exposed to exogenous FFAs (Oleic acid; 0.25 mM). Basal oxygen consumption rates (OCR) were measured. Average values are shown +/- standard errors. Expression of CD36 (B) or LIPG (C) was analysed by qRT-PCR and mean values +/- standard errors are shown. (D, E) *Epas1^EC-KO^* mice and littermate controls lacking Cre (*Epas1^EC-WT^*) were injected with tamoxifen aged 6 weeks and analysed 2 weeks later. *En face* staining of LOSS and HSS regions of the aortic arch using anti-CD36 (D) or anti-LIPG (E) antibodies. Endothelium was co-stained (EC; green) and nuclei detected (DAPI; blue). Each data point represents an animal. Differences between means were analysed using a one-way ANOVA (A), t-test (B, C) or two-wau ANOVA (D, E).

It is notable that the reduction in oxygen consumption in *EPAS1*-silenced cells could be rescued by supplementing cultures with exogenous FFAs (Fig. 6A). This suggests that EPAS1 may regulate the initiation of the fatty acid β-oxidation pathway by controlling the ability of EC to produce or handle FFAs. This was tested by quantifying levels of LIPG and CD36 which act together with ANGPTL4 to co-ordinate FFA transport^37^. RT-PCR analysis showed that *EPAS1* silencing consistently reduced the expression of *LIPG* and *CD36* at the mRNA level (Fig. 6B and Fig. 6C). To confirm the influence of *Epas1* on CD36 and LIPG we performed *en face* staining of the murine aorta. In wild-type mice, CD36 and LIPG were significantly enriched at the inner curvature of the aortic arch (LOSS region) compared to the outer curvature (HSS region) (Fig. 6D and Fig. 6E). The expression of CD36 and LIPG was significantly reduced at LOSS regions in *Epas1^EC-KO^*compared to controls (Fig. 6D and Fig. 6E), indicating that *Epas1* is required for the enrichment of both these molecules at the LOSS region. These data suggest that EPAS1 is required for fatty acid β-oxidation in sheared endothelium by positively regulating the expression of CD36 and LIPG which co-ordinate FFA handling.

Endothelial fatty acid β-oxidation is primarily used to drive dNTP production for DNA repair and proliferation and we therefore hypothesized that EPAS1 may be required for supporting endothelial proliferation via fatty acid metabolism. This was tested by analysis of the murine aorta by *en face* staining of Ki67 (a marker of mitotically-active cells). EC proliferation was significantly reduced in *Epas1^EC-KO^* mice compared to controls (Fig. S9A), indicating the endothelial *Epas1* is required for EC proliferation. Consistently, analysis of cultured PAEC exposed to LOSS revealed that proliferation was significantly reduced by silencing of *EPAS1* (Fig. S9; I vs II). To reconcile the effects of EPAS1 on fatty acid b-oxidation and proliferation we rescued fatty acid metabolism in EPAS1-silenced PAEC by treating them with exogenous FFAs. Metabolic rescue significantly enhanced the frequency of EC proliferation in EPAS1-silenced PAEC (Fig. S9; II vs IV), suggesting that EPAS1 drives proliferation by promoting fatty acid metabolism.

Overall, our data suggest that EPAS1 plays an atheroprotective role at regions exposed to disturbed flow by preserving EC metabolic homeostasis and proliferative capacity.

## DISCUSSION

Our observations indicate a preferential expression of EPAS1 at atheroprone sites exposed to disturbed flow. Under these hemodynamic conditions, the ensuing Laminar Oscillatory Shear Stress (LOSS) promotes atherosclerosis through the activation of multiple pathways, including BMP4-HOXB9^38^, GATA4-TWIST1-SNAIL^39,40^, and NOTCH4-JAG1^29^, collectively increasing inflammation and impeding vascular repair. Surprisingly, in contrast to most LOSS-pathways, our murine model demonstrates that endothelial Epas1 exhibits a protective effect against atherosclerosis. This suggests a potential homeostatic function for EPAS1 expression at atheroprone sites, mitigating the atherogenic process.

A pivotal finding from our research highlights the loss of EPAS1’s homeostatic role during obesity, a global epidemic and major contributor to cardiovascular disease. Our study concludes that obesity leads to the suppression of EPAS1 in arterial endothelial cells, unveiling a novel molecular mechanism contributing to obesity-driven atherosclerosis. Our subsequent observations propose a metabolic regulation of EPAS1 levels, with hypertriglyceridemia potentially suppressing EPAS1 expression. Notably, sulforaphane lowers TGs and rescues EPAS1 expression in obese mice thereby establishing an association between TG and EPAS1 suppression. However, it was important to assess directly if the TG pathway can alter EPAS1 because sulforaphane also has TG-independent effects on endothelium by altering oxidant status^41^. In cell culture conditions EPAS1 expression was reduced by the addition of FFAs which are metabolic byproducts of TGs, suggesting that TG may play a role in EPAS1 suppression. These observations emphasize the role of EPAS1 in obesity-related atherosclerosis.

Further investigation into the mechanism of atheroprotection reveals EPAS1’s requirement for fatty acid β-oxidation in atheroprone endothelium exposed to LOSS. FFAs are generated at the endothelial plasma membrane by hydrolysis of triglycerides via a complex containing LIPG^37^. They are subsequently transported into the cell via CD36. EPAS1 appears to play a role in fatty acid handling, with genetic deletion of Epas1 demonstrating a reduction in LIPG and CD36 protein expression at sites of LOSS. This effect is replicated in PAEC with EPAS1 silencing.

Despite endothelial cells primarily utilizing glycolysis for energy production^42^, we demonstrate that EPAS1-dependent fatty acid metabolism is crucial for proliferation, a process essential for vascular repair. This observation aligns with earlier reports of metabolic dysfunction in Epas1 mutant mice^43,44^ and suggests a potential atheroprotective mechanism through EPAS1-driven endothelial cell proliferation. However, the relationship between endothelial cell turnover and atherosclerosis is intricate, as proliferation is essential for vascular repair but excessive proliferation may contribute to increased permeability to proatherogenic lipoproteins and endothelial activation. We propose that EPAS1 is crucial for maintaining reparative endothelial cells that promote homeostasis, ultimately reducing atherosclerosis.

Our study contrasts the functions of HIF1A and EPAS1 in arterial endothelial cells, with HIF1A promoting atherosclerosis at sites exposed to LOSS^9^. We conclude that these factors have divergent roles in arterial EC, with HIF1A promoting and EPAS1 protecting against atherosclerosis, respectively. Identifying mechanisms that shift the balance of HIF transcription factors to suppress HIF1 and enhance EPAS1 may offer therapeutic opportunities to promote vascular repair and reduce atherosclerosis.

In summary, our findings underscore the preferential expression of EPAS1 at atheroprone sites, where it reduces atherosclerosis by promoting fatty acid metabolism to support endothelial repair. This protective pathway is compromised by obesity, revealing a novel molecular link connecting systemic cardiovascular risk to endothelial cell dysfunction. These findings hold potential implications for the clinical management of obese individuals with cardiovascular risk.

## Acknowledgements

We thank the Adipositazentrum at Limmattal-Hospital, Schlieren for their support in collecting blood samples in patients with obesity.

## Funding

British Heart Foundation RG/19/10/34506 (PCE)

British Heart Foundation Intermediate Fellowship (FS/18/2/33221; JSC)

UKRI Future Leaders Fellowship (MF)

A*STAR Intramural Funding (WH), Strategic Program Fund (BBI; WH) and Central

Research Fund (WH).

Swiss National Science Foundation (PRIMA: PR00P3_179861/1; EO)

Swiss Life Foundation, Switzerland (EO)

Alfred and Annemarie von Sick Grants for Translational and Clinical Research

Cardiology and Oncology (EO)

Heubergstiftung and the Swiss Heart Foundation, Switzerland (EO)

National Institutes of Health grants (R01 HL148239 and R01 HL164577; CLM)

Leducq Foundation Network of Excellence grant ‘PlaqOmics’ (18CVD02; CLM).

A*STAR Research Attachment Program (DP).

Academy of Medical Sciences Springboard Award (SBF005/1064; SPA).

PGC 2018-094025-B-I00 funded by MCIN/AEI/10.13039/501100011033 (GV)

Fondo Europeo de Desarrollo Regional (FFEDER) A way of making Europe (GV).

## Declaration

Authors declare that they have no competing interests.

